# Acoustic and Neurophysiological Aspects of Lombard Effect

**DOI:** 10.1101/2022.09.30.510383

**Authors:** Christian Castro, Lucia Z Rivera, Pavel Prado, Jhosmary Cuadros, Juan Pablo Cortés, Alejandro Weinstein, Victor Espinoza, Matías Zañartu

## Abstract

**Purpose:** This study aims to describe variations in acoustic and electroencephalography measures when speaking in the presence of background noise (Lombard effect) in participants with typical voice and normal hearing.

**Method:** Twenty-one participants with typical voices and normal hearing uttered simple vocal tasks in three sequential background conditions: Baseline (in quiet), Lombard (in noise), and Recovery (five minutes after removing the noise). Acoustic and electroencephalography signals were recorded in all conditions. The noise used in the Lombard condition consisted of speech-shaped noise at 80 dB SPL sent by headphones. Acoustic measure, and ERP responses were analyzed.

**Results:** During the Lombard condition, the participants increased the intensity of their voice, accompanied by an increase in CPP, and a decrease in H1-H2. The cortical response was characterized by the increased N1-P2 complex amplitude of the ERP elicited by the subject’s own vocalizations in noise, The source localization showed neural activities in frontal and temporal cortical regions.

**Conclusions:** The variation in acoustic measures due to the Lombard Effect could be modulated by temporal, and cortical regions.

## Introduction

The voice production depends on the acoustic context, reflected when individuals talk in noise environments. Noise affects the auditory feedback of their own voice, generating obstacles in communication. Thus, Speakers involuntarily increase the vocal intensity to ensure the message reaches the target audience, a process that has been referred to as Lombard effect (LE) (Lombard, 1911 In addition, to variations in intensity, LE includes (i) an increase in the fundamental frequency of voice (Stowe & Golob, 2013); (ii) variations in the spectral distribution of acoustic energy of speech; (iii) increment in the regions of second and third formants (Lu & Cooke, 2008) and (iv) the reduction in the length of syllables and vowels (Gardnier, M. et al, 2011). Typically, the LE occurs when the intensity of the background noise exceeds 43 dB(A) (Bottalico et al., 2017). When noise intensity is higher than 55 dB SPL, vocal intensity increases 0.38 dB per dB of background noise (Korn, 1954; Stowe & Golob, 2013). In addition, the LE induced by speech noise (the spectral composition of the background noise resembles that of the speech) is more pronounced than that evoked by background noise without speech frequency components (Garnier, M. et al., 2010; Stowe & Golob, 2013). LE acts as an adaptive communication mechanism and depends on the linguistic context of the message (Patel, R & Schell, K. W., 2008), the type and intention of the communicative interaction among speakers and the audience (Garnier, M. et al., 2010) and the spectral composition of the sound masker (Stowe & Golob, 2013; Meekings et al., 2016).

### Auditory-Motor loop in speech production

The areas involved in the control of phonation are auditory and somatosensory cortex, in the framework of internal model for the sensorimotor integration (Wolpert, 1995, Houde, 2011). During speech production, the supplementary motor area sends simultaneously a signal to motor control regions for producing the desired target sound and a predictive inhibitory signal to auditory, and somatosensory regions (feedforward or “copy efference”). When the sound is produced, the ascending auditory system processes the “current auditory feedback”, which is compared with the predictive feedforward signal. If the auditory feedback matches its prediction, the activity of the auditory cortex in response to one’s own voice, decreases; and the auditory feedback is ignored. On the contrary, when any perturbation of auditory feedback was produced, generates a mismatch between the feedforward and auditory feedback and triggering an error signal. The error signal sends corrective commands to speech muscles for adjustment of the voice, and speech production. Moreover, if the same type of error persists throughout many speech repetitions, then the original speech motor plans might be updated, resulting in long-term “adaptation.” (Behroozmand,2011; Guenther & Hickok, 2015; Houde & Chang, 2015). Thus, during the phonation in a noisy environment, the perception of their own voice is impaired. In detail, when predictive commands (feedforward) do not match with the information of the auditory feedback, an error signal is generated. The involuntary increase in the intensity of voice (due to the masking noise), may act a correction responding to the error signal; generated by the mismatch between feedforward and auditory feedback (Meeking 2016, Meeking 2021).

### Neural basis of Lombard Effect

Based on the model of Dual-Network Model for Vocal–Motor Production, neurological studies propose that subcortical and cortical brain areas would be involved when the talkers speak in noisy environments. Lou, and cols propose the LE is elicited by a subcortical network, which may be modulated by cortical brain areas in mammals or homologous higher level brain structures in vertebrates, such as the pallium in birds (Lou, 2018).

Recent studies using functional magnetic resonance (fMRI) in humans, explored the role of the cortical component of LE. The LE increases the activation of the superior temporal gyrus (STG), and middle temporal gyrus (MTG) during whispered speech and masking noise (Zheng, 2010). Christofelss proposes that when using different levels of masking noise, greater cortical activation in left and right STG is generated. Yet, he explicitly instructs participants to avoid any kind of compensation to the perturbation (masking noise); So that, the restriction of the compensatory effect might limit the cortical activation in LE. (Christofelss (2011). Meekings (2016) find many numbers of subcortical, and cortical areas measured by BOLD signals such as the thalamus, right pallidum, right and left insula, cerebellum lobule, right inferior frontal gyrus, left postcentral gyrus, and bilateral activation in superior temporal gyrus (STG) when subjects speak under noisy condition compared to speaking in quiet. On the contrary, fMRI main studies focus the analysis on the STG as a region of interest (ROI); due to STG acts as an auditory error monitor and might sends a corrective signal to the larynx muscles during auditory masking on the speaking in noise (Meeking, 2021).

The use of fMRI provides useful information about the localization of the cortical areas; Yet, this procedure has limitations in the temporal analysis and the dynamics of the neural network involved in the auditory feedback error signal, and the compensatory response triggered by the masking noise. The variation of BOLD signals in the fMRI paradigm was measured immediately after the speaking in noise, but not during the LE. The temporary limitation of the fMRI might be a bias in the measurement of the LE. Thus, the use of electroencephalography (EEG) could be useful for extending the study of the LE to the analysis of temporal neural response, and the dynamic among the different cortical areas which modulate the compensatory response on the speaking in noise.

#### Aims and hypotheses

The main goal of this study was to explore the neural temporal response and the dynamic of cortical areas involved in the modulation response due to speaking in noise (The Lombard Effect). Thus, the purpose of this study was to compare the voice production and electrophysiological variations among different acoustic backgrounds in individuals with healthy voices. We proposed three different acoustic scenarios for the speakers; (1) In quiet environments, (2) under the Lombard Effect (noisy environment), and (3) five minutes or remove the noise (in quiet). It is hypothesized that when the participants speak in noise, involuntarily increase the intensity of their voice (LE), these acoustic variations would be related to an increase in N1-P2 complex on event-related potentials response (ERP). In addition, we expected an increase in the electrical activity of auditory and somatosensory cortical regions when the participants speak in noise. On the contrary, when the participants speak again in quiet environments (when removing the noise) we hypothesized that a decrease in the intensity of their voice, accompanied by a decrease in N1-P2 complex, and a decrease in the electrical activity of auditory and somatosensory cortex regions.

## Methods

### Participants

Twenty-one volunteers were recruited for this study (mean age = 27.9, SD= 3.6). All participants were assessed by a speech-language pathologist based on case history, clinical evaluation, aerodynamic and acoustic measures of vocal function, and a Consensus Auditory-Perceptual Evaluation of Voice (CAPE-V) (Kempster et al, 2009) The participants showed typical voice with no sign of voice disorder. In addition, they did not report history of speech, language, or hearing disorders. Participants passed a pure-tone hearing screening, which consisted of positive responses to air-conduction stimuli in both ears at 20 dB HL at octave frequencies between 250 Hz and 8000 Hz using a clinical audiometer (Model AD629, Interacoustics A/S, Middelfart, Denmark).

Volunteers provided written consent in accordance with the experimental protocol. The consent was approved by the Research and Ethics Committee of the Faculty of Medicine, Universidad de Valparaíso, Chile; in concordance with the assessment statement code 52015 and in compliance with the national guidelines for research with human subjects and the Declaration of Helsinki.

### Experimental design

The experiment was composed of three sequential acoustic background conditions: Baseline (in quiet), Lombard (in noise), and Recovery (in quiet after five minutes of rest). In each condition, participants were asked to utter eighty syllables at a comfortable pitch and loudness. The syllables consisted in /pa/, /da/, /ta/, and /ba/ presented in a random order. The duration of the vocalization (3 seconds) and pace of the syllable pronunciation (a syllable every six seconds) were controlled by visual cues displayed on a screen (Figure 1). Participants were instructed to speak at a comfortable loudness level while they received the auditory feedback of their own voice through headphones. No specific loudness or pitch was targeted so that the Lombard effect may be appropriately assessed. The voice and EEG signals were recorded in each condition. Using vowels and syllables allowed to obtain reliable and reproducible ERP responses. The limitations and implications of using relatively short vocalizations will be addressed in the discussion section.

**Figure 1.**
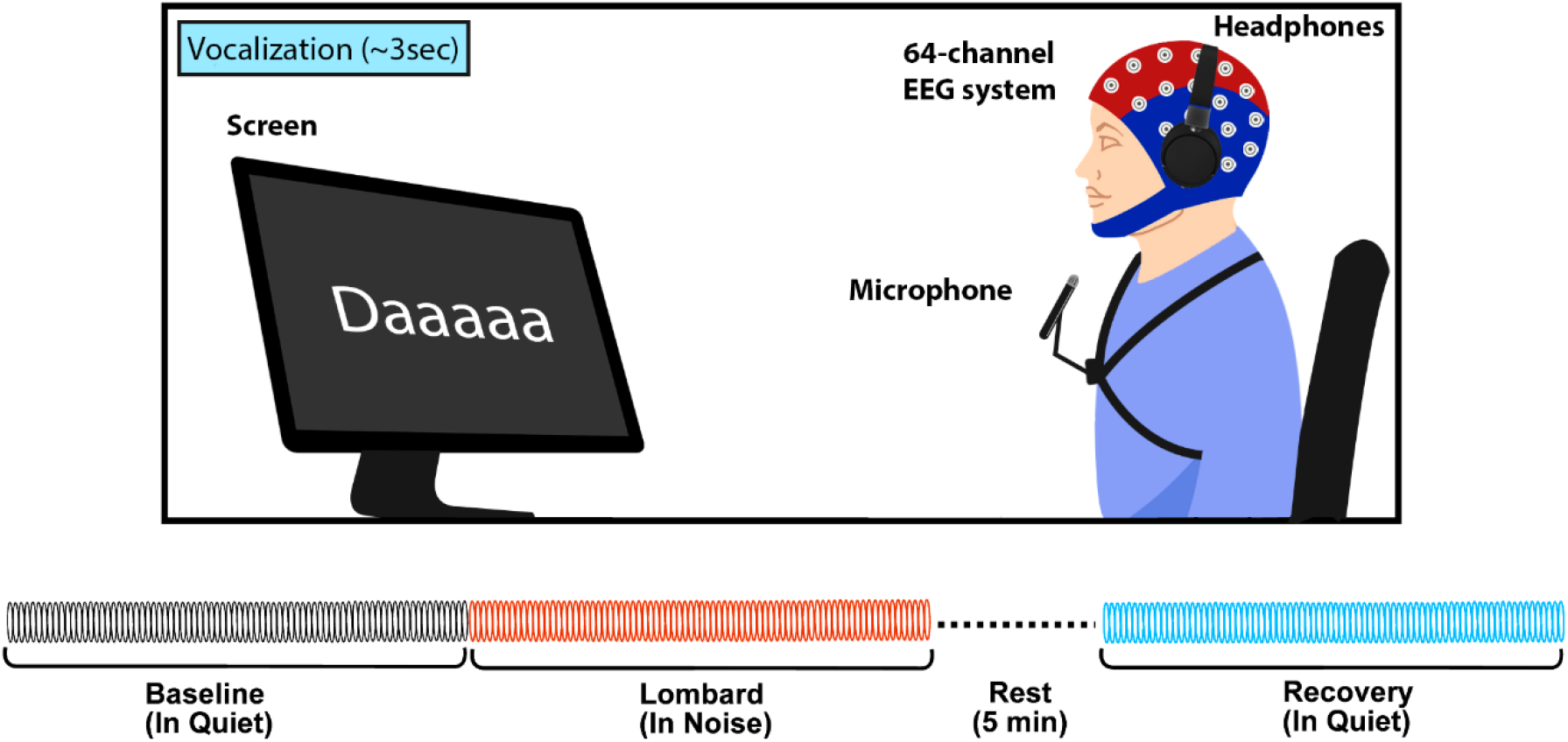
An overview of the experimental design is shown. The experiment involved four conditions: Baseline (in quiet), Lombard (in noise), and Recovery (in quiet after five minutes of rest).

In lombard condition, speech-shaped noise was generated by a clinical audiometer (Model AD629, Interacoustics A/S, Middelfart, Denmark) and presented through the headphones. The noise profile used for triggering the Lombard effect was 80 dB SPL, with equal energy between 125 Hz and 1000 Hz (octave band) followed by an energy decay of 12dB/octave until 6000 Hz. The noise profile is similar to the weighted noise used in previous studies (Junqua, 1993; Van Summers et al., 1988; Lu & Cooke, 2008), aim at inducing a robust Lombard effect while avoiding hearing discomfort and vocal/auditory fatigue (Alghamdi et al., 2018). Due to the closed-ear design of the headphones, a certain degree of attenuation of the voices of the participants could have occurred. However, use of these headphones has previously shown no significant effect on speech acoustics (Lu & Cooke, 2008).

The recovery condition was included to explore the potential persistence of the Lombard effect after the speaking under noise. A five-minute rest between the Lombard and Recovery conditions was selected for allowing participants to recover from possible vocal loading effects, as suggested in previous studies (Fujiki & Sivasankar, 2017; Xue et al., 2019).

### Acoustic measures

The acoustic signal was obtained using a microphone (B&K, model 4961; Nærum, Denmark) located in front of the participant at 15 cm from the lips at a 45-degree offset in the axial direction and amplified by a B&K 1705 signal conditioner. The acoustic signal was calibrated to physical units of dB SPL (dB re 20 μPa) using a Larson Davis calibrator (model CAL200, Depew, NY, USA). Next, signals were sampled at 20 kHz with 16-bit quantization and low-pass filtered (3 dB cutoff frequency of 8 kHz) using a National Instrument DAQ model USB-6363 BNC. From acoustic signal the vocal SPL was computed for each vocalizations using a window size ~200 ms. For testing our hypotheses, repeated measures analyses of variance (ANOVA) were performed to analyze the changes of the acoustic SPL measures. A Post hoc tests using Bonferroni’s correction was realized to test the statistical significance between condition (α = 0.05).

### EEG data acquisition and analysis

The EEG signals were recorded using a BioSemi ActiveTwo system with 64 active electrodes, at a sampling rate of 4096 Hz. The location of each active Ag/AgCl electrode was according to the standard 10–20 montage. Then synchronized the EEG data with acoustic recordings.

The continuous EEG data was pre-processed offline using Brain Vision Analyzer 2.0^®^ software (Brain Products GmbH, Munich, Germany) following standard procedures for EEG (Delorme & Makeig, 2004), to calculate ERPs responses time-locked. Recordings were first band-pass filtered among 0.1-30 Hz with a zero-phase shift Butterworth filter of order 8. Then, down sample rate to 512 Hz. Ocular, muscular and line noise artifacts were corrected and removed by visual inspection with the Independent Component Analysis (ICA) method. (following Chaumon et al., 2015). Data were re-referenced to the average of all channels and epochs were segmented −200 to 500 ms around to the onset of vocalization. Epochs were rejected if the amplitudes exceeded a maximum voltage of ± 50 μV. In the epochs average, ICA was used again to identify the ERP components based on their latency, topography, and polarity. Baseline correction was performed to the prestimulus period.

The statistical analysis of the ERP responses was performed in a 2-point average window around the peak of the N1 and P2 components with respect to the onset of vocalization. Then, subtraction of these values was performed to analyze the variation in the N1-P2 complex. A repeated measures ANOVA test was used including the conditions of N1-P2 baseline, N1-P2 lombard and N1-P2 recovery, for the electrodes grouped in a position of interest (O1, O2, Oz, P1, P10, P2, P3, P4, P5, P6, P7, P8, P9, PO3, PO4, PO7, PO8, POz, Pz).

### ERP source localization

Source localization was estimated using the Bayesian Model Averaging (BMA); BMA calculates the brain sources considering the anatomical constraints to solve the EEG/MEG inverse problem. Thus, it favors brain regions and penalizes those in relation to probabilities of contributing to the generation of the data. Moreover, allows for the estimation of deep EEG generators. (For a review, refers to Trujillo-Barreto et al., 2003).

Cortical activation maps were calculated for each scalp voltage distribution based on the latencies of the N1-P2 components of the ERP. A single time window between ±10 ms around the maximum amplitude values of N1-P2 was taken for the baseline, lombard, and recovery conditions. The statistical comparison of the ERP cortical activity maps was performed in pairs, using a T test (α = 0.05), considering Lombard v/s Baseline, Lombard v/s Recovery and Recovery v/s Baseline contrasts. Multiple voxel-by-voxel comparisons were corrected for by implementing a 5000 nonparametric permutation test.

## Results and discussion

We assessed the variations in the acoustic measures of voice, and the variation in neurophysiological activity under three conditions: speaking in a quiet environment (Baseline condition), speaking under masking noise (Lombard condition), and speaking after 5 minutes of rest in a quiet environment (Recovery condition). These conditions were selected to explore how speaking in noise affects phonation and to describe the potential neurophysiological basis underlying the LE.

### Accoustic measures

When participants spoke under masking noise (Figure 2, a), the LE was evident. Thus, their voice intensity (SPL) increased from the Baseline condition to the Lombard condition; Then, decreased in the recovery condition (in quiet), five minutes after the speech noise was removed with a significant difference in the SPL values among conditions (F=44.210, p<0.001). In detail, a Post hoc test using Bonferroni’s correction referring a significant difference between Lombard and baseline (p<0.001), Lombard and recovery (p<0.001) condition on behavioral responses during vocal production. No significant main effect was present between baseline and recovery conditions.

**Figure 2.**
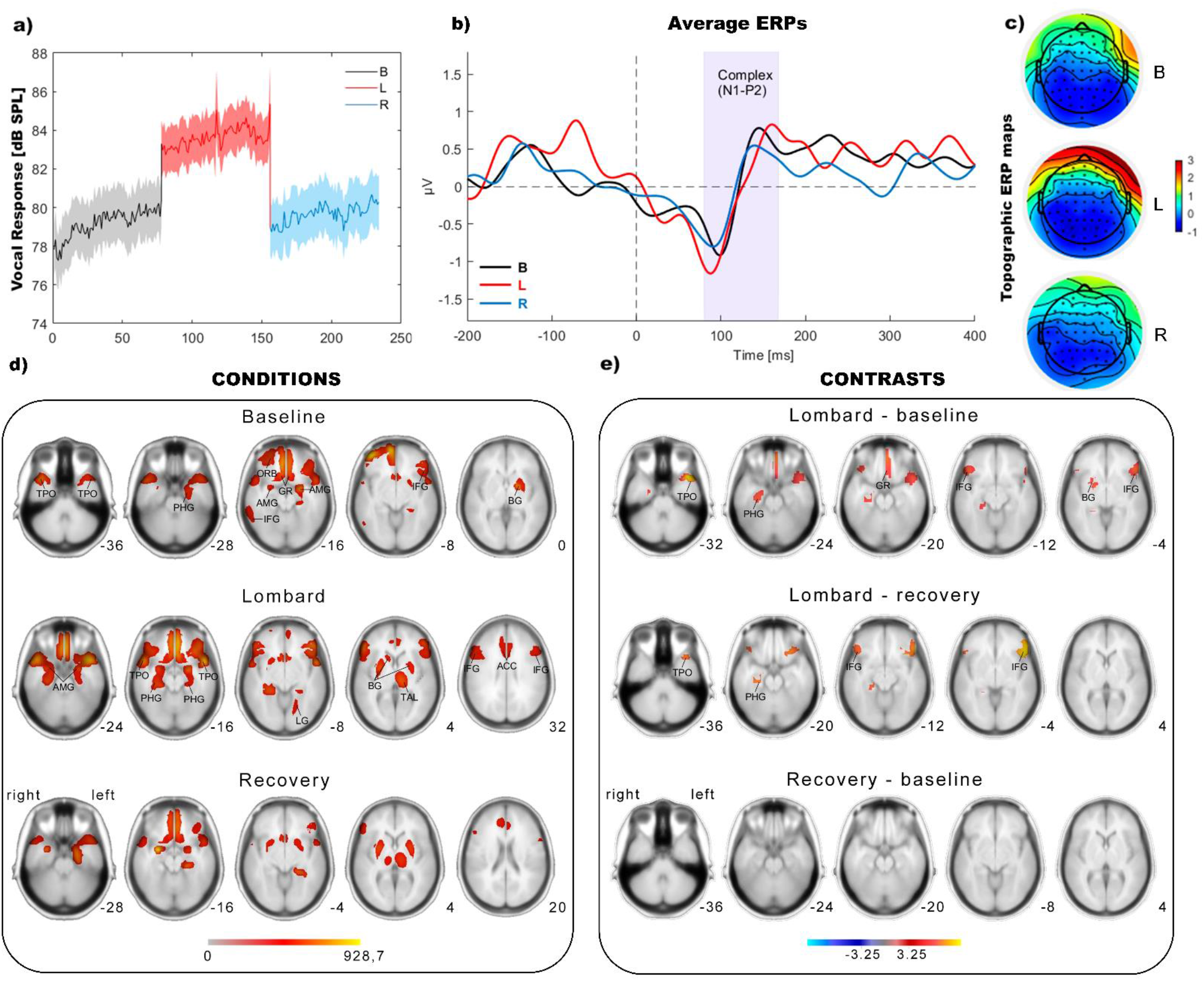
**a)** Results of the behavioral vocal response analysis of the acoustic SPL measures: profile of the grand-average (n=21) during Baseline, Lombard (in noise), and Recovery conditions. **b)** Grand mean ERPs in all subjects at a pooling of electrodes (average of O1, O2, Oz, P1, P10, P2, P3, P4, P5, P6, P7, P8, P9, PO3, PO4, PO7, PO8, POz, and Pz). **c)** Topographic distribution maps for each condition on a scalp map of 64 EEG electrodes. **d)** Source localization for each condition.**e)** Representation of the statistical analysis of the source localization for the contrasts between lombard-baseline, lombard-recovery and recovery-baseline.

Our results showed that participants generated a compensatory response to masking noise; with an increase in their SPL, compared to their Baseline condition (in a quiet environment). The participants with typical voices increased their SPL by a mean of 5.2 dB when speaking in noise. This variation is concordant with previous studies of the LE; in which, the experimental condition used masking noise (Meekings et al., 2016; Stowe & Golob, 2013; Garnier et al., 2010).

The variation in SPL of the voice of participants due to the speaking in noise was accompanied by significant variation in H1-H2 (F=44.210, p<0.001), and CPP (F=44.210, p<0.001) among conditions. In detail, Post hoc tests using Bonferroni’s correction referring a significant difference between Lombard and baseline (p<0.001), Lombard and recovery (p<0.001) condition on behavioral responses during vocal production for H1-H2. In the same direction, for CPP Post hoc tests using Bonferroni’s correction referring to a significant difference between Lombard and baseline (p<0.001), Lombard and recovery (p<0.001) condition. No significant main effect was present between baseline and recovery conditions for both measures (H1-H2 and CPP).

The decrease in H1-H2 under masking noise conditions could be attributed to an increase in the velocity of vibration on the vocal fold due to the LE. On the other hand, speaking in noise modify the spectral component of speech (Gardnier, 2013), this modification could impact variation on voice quality expressed in the increase of CPP.

### ERP responses and source localization

The ERPs for each condition and the topographic distribution maps are present at the top and middle of Figure 2,b. The results of the statistical analysis indicated a significant effect among the experimental conditions (F=7.943, p=0.001). Post Hoc tests using the Bonferroni correction revealed that the amplitude of the N1-P2 complex was significantly higher for the lombard condition compared to baseline (p=0.009) and recovery (p=0.002). However, the amplitude of the N1-P2 complex did not differ significantly between Baseline and Recovery conditions (p= 0.62).

Source localization analyzes of the N1-P2 complex of the ERP elicited by one’s own voices revealed distributed brain activation within the frontal, limbic and temporal lobe (Figure 2, d). The topographic distribution of the N1-P2 generators was consistent across conditions (Baseline, Lombard, and Recovery), and included brain areas that are i) crucial for speech comprehension, ii) relevant for the processing of the auditory feedback of one’s own voices, and iii) that are activated due to listening efforts in noisy environments. In the frontal lobe, the N1-P2 generators included the orbitofrontal gyrus (ORB), the gyrus rectus, and the inferior frontal gyrus (IFG). In the limbic lobe, sources of the ERP were estimated in the hippocampus/parahippocampal gyrus (PHG), and the amygdala (AMG). Lastly, temporal areas associated with the N1-P2 generation included the temporal pole (TPO), and the inferior frontal gyrus (ITG).

In comparison with the brain activity elicited by the auditory feedback of one’s own voices in quite (baseline), self-monitoring of speech in noise (Lombard) was associated with higher activations in the IFG, the GR, the PHG, and the basal ganglia (BG) (voxel wise t-test: p ≤ 0.05, corrected for multiple comparison using permutations test, 5000 randomizations) (Figure 2, e). The same areas, with the exception of the GR and the BG, also displayed hyperactivation during the Lombard effect, in relation to the brain activity estimated in the Recovery condition (voxel-wise t-test: p ≤ 0.05, corrected for multiple comparison using permutations test, 5000 randomizations) (Figure 2, e). In neither of these contrasts, cortical hypoactivation was associated with the self-monitoring of speech in noise (voxel-wise t-test: p ≥ 0.05). Likewise, the brain activation did not significantly differ between the Baseline and the Recovery conditions (voxel-wise t-test: p ≥ 0.05) (Figure 2, e).

The LE was still reliably elicited even though the communicative environment was greatly simplified/limited (production of short monosyllables and absence of visual contact with a communication partner). In fact, the variation of SPL we observed is similar to that obtained in studies using running speech or linguistic corpus (Stowe & Golob, 2013; Patel & Schell, 2008; Garnier et al., 2010). This suggests that a minimum communicative intention is needed to elicit the LE, and that the quality and complexity of the message is a secondary element in this adaptive behavior. This is supported by evolutionary studies demonstrating the widespread distribution of the Lombard effect in vertebrates, present in large numbers of species that do not have complex oral communication (for a review, Luo et al., 2018).

## Conclusion

The Lombard effect generates variations in acoustic measures linked with an increase in the amplitude and velocity of vocal folds vibration. This variation could be modulated by temporal and frontal brain activation.

## Acknowledgment

This research was supported by Agencia Nacional de Investigación y Desarrollo (ANID) through grants FONDECYT 1191369, BASAL FB0008, ANILLO ANID/ACT210053 and the National Institutes of Health (NIH) National Institute on Deafness and Other Communication Disorders grant P50 DC015446. The content is solely the responsibility of the authors and does not necessarily represent the official views of the National Institutes of Health.

## Notes

### Competing Interest Statement

The authors have declared no competing interest.

